# Intramuscular adrenaline administration does not improve a survival period on rats with crush syndrome despite short-term hemodynamic support and renal protection

**DOI:** 10.64898/2026.04.30.722096

**Authors:** Isamu Murata, Yoshiaki Miyamoto, Jun Kobayashi

## Abstract

Crush syndrome (CS) is a serious medical condition characterized by damage to the muscle cells due to pressure and is associated with high mortality, even in patients receiving fluid therapy. We focused on adrenaline (Adr), a standard medication administered by medical teams dispatched during disasters. Adr is readily available for use in disaster scenarios owing to its inclusion in standard emergency kits. The effectiveness of Adr in the treatment of CS remains a subject of ongoing debate. This study aimed to evaluate the impact of Adr on acute complications, such as heart failure, shock, and renal failure, and explore whether its influence on inflammatory pathways is correlated with improved survival in rats with CS. The CS model involved subjecting anesthetized rats to bilateral hindlimb compression using a rubber tourniquet for 5 h. Subsequently, the rats were randomly divided into eight groups. Under continuous monitoring and recording of the arterial blood pressure, blood and tissue samples were collected for biochemical analyses at designated time points before and after reperfusion. The survival rate, vital signs, and blood gas parameters were higher in the CS group than in the sham group. They were improved in the Adr-treated group (0.01 or 0.01 mg/kg), which was not significantly different from that in the CS group, despite the improvement in shock and kidney dysfunction. In conclusion, intramuscular Adr provides immediate hemodynamic stabilization and renal protection during the early stages of CS. However, its use requires careful dose titration; low doses may promote the systemic release of lethal toxins, whereas high doses may worsen metabolic acidosis. These findings highlight the importance of combining Adr with other therapies, such as fluid resuscitation, to manage systemic toxemia inherent in CS.

## Introduction

Crush syndrome (CS) is a life-threatening traumatic condition that is frequently reported during large-scale earthquakes. CS is a systemic condition resulting from skeletal muscle cell injury caused by compression or crushing. The pathogenesis of CS progresses as follows: muscle destruction and ischemia–reperfusion injury caused by limb compression leading to the release of potassium (K^+^), myoglobin (MB), and pro-inflammatory factors from damaged muscle cells into the systemic circulation (1-4). This process triggers hypovolemic shock, heart failure due to hyperkalemia, myoglobinuric acute kidney injury, and systemic inflammatory response syndrome. Although approximately 20% of fatalities occur in the ultra-early phase due to cardiac arrest or similar causes (5), CS remains a high-mortality trauma, in which deaths from cytokine storm-related inflammatory diseases simultaneously occur over the medium-to-long term. The treatment strategy for CS focuses on managing hypovolemic shock and hyperkalemia and preventing renal failure. Current guidelines recommend aggressive fluid resuscitation during early interventions. However, in large-scale disaster situations where medical resources are severely limited, sustaining high-volume fluid therapy, which primarily addresses hyper-acute symptoms, is often challenging. Furthermore, the onset of symptoms related to potentially progressive inflammatory responses requires even more medical supplies, thus complicating the situation.

We focused on adrenaline (Adr), a standard medication administered by medical teams dispatched during disasters. Adr is readily available for use in disaster scenarios owing to its inclusion in standard emergency kits. It acts as a β-receptor agonist, increasing cardiac output and heart rate (HR), and as an α-receptor agonist, inducing vasoconstriction. Suppressing blood flow through vasoconstriction in crushed muscles can delay the systemic circulation of substances, such as K^+^, thereby mitigating the progression of heart and renal failure and inflammatory responses (6). The adrenergic effect is expected to suppress the systemic release of K^+^ from the injured sites and promote cellular K^+^ uptake. However, some reports have suggested that Adr may conversely promote inflammation in the medium-to-long term (7). Therefore, the effectiveness of Adr for the treatment of CS remains a subject of ongoing debate. Thus, this study aimed to evaluate the impact of Adr on acute complications, such as heart failure, shock, and renal failure, and explore whether its influence on inflammatory pathways is correlated with improved survival in rats with CS.

## Materials and methods

### Animal and crush syndrome (CS) model rat

Animal CS model. Male Wistar rats weighing 250-300 g were obtained from Japan SLC (Shizuoka, Japan) and housed in a room maintained at a temperature of 23°C ±3 °C and relative humidity of 55±15% with a 12/12-h light/dark cycle and free access to food and water. All animal experiments were approved by the Institutional Animal Care and Use Committee of Josai University (approval nos.: H29030 and JU18031). Anesthesia was induced and maintained using 2% isoflurane. The body temperature (BT) was maintained throughout the experiment using a heating pad. The CS model was established, as previously described (4). Briefly, a rubber tourniquet was applied to both hind limbs (HLs) of each rat, which was wrapped five times around a 2.0 kg metal cylinder, and the end of the band was glued. Immediately before compression for 5 h, the tourniquet was removed from the limbs by cutting the band (i.e., reperfusion for 0 h).

### Experimental design

The Adr used in this study was Bosmin® Injection 1 mg (Daiichi Sankyo Co., Ltd.). Adr doses of 0.01 and 0.1 mg/kg were prepared, each diluted in 100 μL of saline. The anesthetized rats were randomly assigned to the following seven groups

#### Sham and S-AL0.1 groups

The rats underwent 5 h of anesthesia induction without compression, followed by an injection of either 100 μL saline (vehicle) or 0.01 mg/kg Adr into the anterior limb (AL) (injection site: right triceps brachii).

#### CS, C-AL0.01, C-AL0.1, C-HL0.01, and C-HL0.1 groups

CS was induced using a rubber band for compression. Immediately before decompression, 100 μL of saline (vehicle) or 0.01 or 0.1 mg/kg Adr was injected into either the AL or compressed area of the HL.

#### CL group

To evaluate the effect on blood flow, the CL (clamping) group underwent the same compression procedure as the CS group. Immediately before decompression, the femoral arteries and veins in both inguinal regions were ligated with silk sutures (Natsume Seisakusho Co., Ltd.), and the skin was closed using polyamide sutures (Matsuda Ika Kogyo Co., Ltd.).

#### Experiment 1 (survival rate)

To examine the survival period, the rats were randomly divided into eight groups (n=15 each): (i) sham, (ii) S-AL0.1, (iii) CL, (iv) CS, (v) C-AL0.01, (vi) C-AL-0.1, (vii) C-HL0.01, and (viii) C-HL0.1. All the rats used in the experiments were euthanized (confirmation by pupillary reflex to light) by sodium pentobarbital overdose (100 mg/kg body weight, intravenous administration) at 48 h after all measurements were recorded.

#### Experimental design 2 (vital signs, blood gas levels, and biochemical parameters)

To examine the changes in blood pressure and blood gas parameters, the rats were randomly divided into the following eight groups (n=6 each): (i) sham, (ii) S-AL0.1, (iii) CL, (iv) CS, (v) C-AL0.01, (vi) C-AL-0.1, (vii) C-HL0.01, and (viii) C-HL0.1. Sequential sampling was performed at 0.5, 1, 3, 6, and 24 h after reperfusion. Regarding vital signs, blood gas levels, and biochemical parameters, the following vital signs were recorded using a PowerLab data acquisition system (AD Instruments, Nagoya, Japan): systolic blood pressure (SBP), mean blood pressure, diastolic blood pressure, HR, and BT. One carotid artery was cannulated using a polyethylene catheter (PE-50 tubing) and connected to a pressure transducer. Arterial blood samples were obtained from each group at 0.5, 1, 3, 6, 12, and 24 h after reperfusion using the carotid artery catheter. The arterial levels of pH, arterial partial pressure of carbon dioxide, partial pressure of oxygen, lactate, total carbon dioxide, bicarbonate, oxygen saturation, base excess (BE), sodium ion, K^+^, chloride, glucose, blood urea nitrogen (BUN), hematocrit, hemoglobin, and anion gap were analyzed using an i-STAT300F blood gas analyzer with CG4+ and EC8+ cartridges (FUSO Pharmaceutical Industries, Osaka, Japan). Arterial blood was collected from the jugular vein and centrifuged to measure plasma creatine phosphokinase (CPK) using a Creatine Kinase Assay Kit (BioAssay Systems Co., CA, USA), Mb using a solid-phase enzyme-linked immunosorbent assay (Rat Myoglobin ELISA; Life Diagnostics, West Chester, PA, USA), and creatinine using a Creatinine (serum) Colorimetric Assay Kit (Cayman Chemical, USA). To assess kidney function, the bladder was cannulated parallel to the jugular vein using PE-50 tubing. Urine samples were obtained 1 h before and immediately after decompression (0 h). Kidney function was determined based on the urine volume and urine pH (Pretest 5bII; Wako Pure Chemical Industries). Samplings for plasma interleukin (IL)-6 and atrial natriuretic peptide (ANP) were performed at 24 h after reperfusion, IL-6 level was measured using the Quantikine® ELISA kit (RandD Systems), and plasma ANP level was measured using the Rat ANP ELISA Kit (NPPA) (Abcam). All the rats used in the experiments were euthanized (confirmation by pupillary reflex to light) by sodium pentobarbital overdose (100 mg/kg body weight, intravenous administration) at each sampling time.

#### Experimental design 3 (blood perfusion in the crushed hind limb)

Blood perfusion (BPF) in the crushed HL was evaluated in the rats divided into the following eight experimental groups (n=6 each): (i) sham, (ii) S-AL0.1, (iii) CL, (iv) CS, (v) C-AL0.01, (vi) C-AL-0.1, vii) C-HL0.01, and (viii) C-HL0.1. Sequential measurements were performed at 0.5, 1, 3, 6, and 24 h after reperfusion. Regarding BPF parameters, the BPF of the muscles and blood vessels was measured using PeriScan PIM3 (PERIMED, Stockholm, Sweden) in experimental design 3. BPF was assessed under the following conditions: distance to the hind limb, 10-15 cm; measurement area, 10×10 cm; environmental temperature, 23±5 °C; and relative humidity, 50±5% (6). All the rats used in the experiments were euthanized (confirmation by pupillary reflex to light) by sodium pentobarbital overdose (100 mg/kg body weight, intravenous administration) at 24 h after all measurements were recorded.

### Statistical analyses

Results are expressed as means±standard errors of the mean. Differences between the groups were assessed by analysis of variance with Tukey’s honest significant difference test or the Tukey–Kramer test. Survival curves were generated using the Kaplan-Meier method, and survival was compared using the log-rank test. P<0.05 was considered statistically significant (Statcel 2, 2nd ed., OMS Publishing Inc., Saitama, Japan).

## Results

### The effects of adrenaline on CS rat viability

Survival rates are shown in Fig. 1. The survival rates of rats in the CS group were 93%, 87%, 53%, 33%, and 30% at 1, 3, 6, 24, and 48 h after reperfusion, respectively. Three hours after reperfusion, the survival rates in the C-AL0.01, C-AL0.1, C-HL0.01, and C-HL0.1 groups were similar to those in the CS group. Until 6 h after reperfusion, the survival rates in the C-AL0.01, C-HL0.01, and C-HL0.1 groups were similar than those in the CS group. However, the survival rate in the C-AL0.1 group was significantly lower than that in the CS group. At 48 h after reperfusion, the survival rates were not significantly different from those in the CS group. In the experimental period, the rats in the sham and S-AL0.1 groups did not die. Furthermore, the survival rate in the CL group was significantly higher than that in the CS group at 48 h after reperfusion.

**Figure 1.**
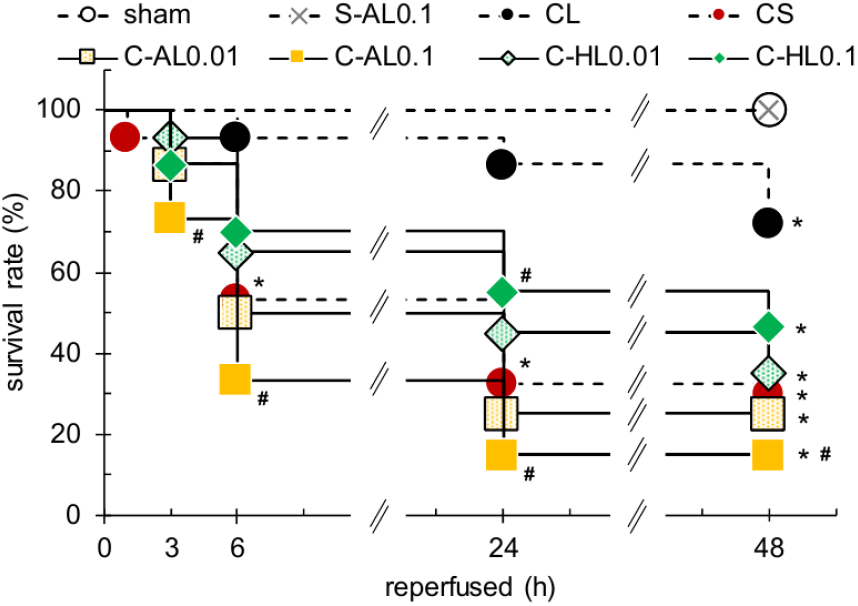
The impact of intramuscular adrenaline of varying durations on CS rat viability. White circle, sham; black cross, S-AL0.1; black circle, CL; red circle, CS; yellow dot square, C-AL0.01; yellow square, C-AL0.1; green dot diamond, C-HL0.01; and green diamond, C-HL0.1. Survival curves are shown using the Kaplan–Meier method (n=15). ^*^p<0.05 vs. sham group, ^#^p<0.05 vs. CS group, ^†^p<0.05 vs. C-AL-0.01, ^♭^p<0.05 vs. C-AL-0.1, ^§^p<0.05 vs. C-HL-0.01 (log-rank test).

### Effects of intramuscular adrenaline on biochemical markers

CPK levels were measured to evaluate the effectiveness against K^+^-induced heart failure due to muscle injury (Fig. 2). The CS group had significantly higher plasma CPK and K^+^ levels than the sham group at all experimental periods (CPK, 15.3×10^3^±1.5×10^3^ vs. 0.35×10^3^±0.08×10^3^ IU/L, p<0.05; K^+^, 7.9±0.3 vs. 3.9±0.2 mmol/L, p<0.05, reperfused at 24 h). In terms of CPK levels, the Adr-treated groups showed no significant difference with the CS group. In terms of the plasma K^+^ levels, the C-AL0.01 and C-AL0.1 groups had significantly higher plasma K^+^ levels than the CS group at 3–24 h after reperfusion. In contrast, the C-HL0.01 and C-HL0.1 groups had significantly lower plasma K^+^ levels than the CS group 24 h after reperfusion.

**Figure 2.**
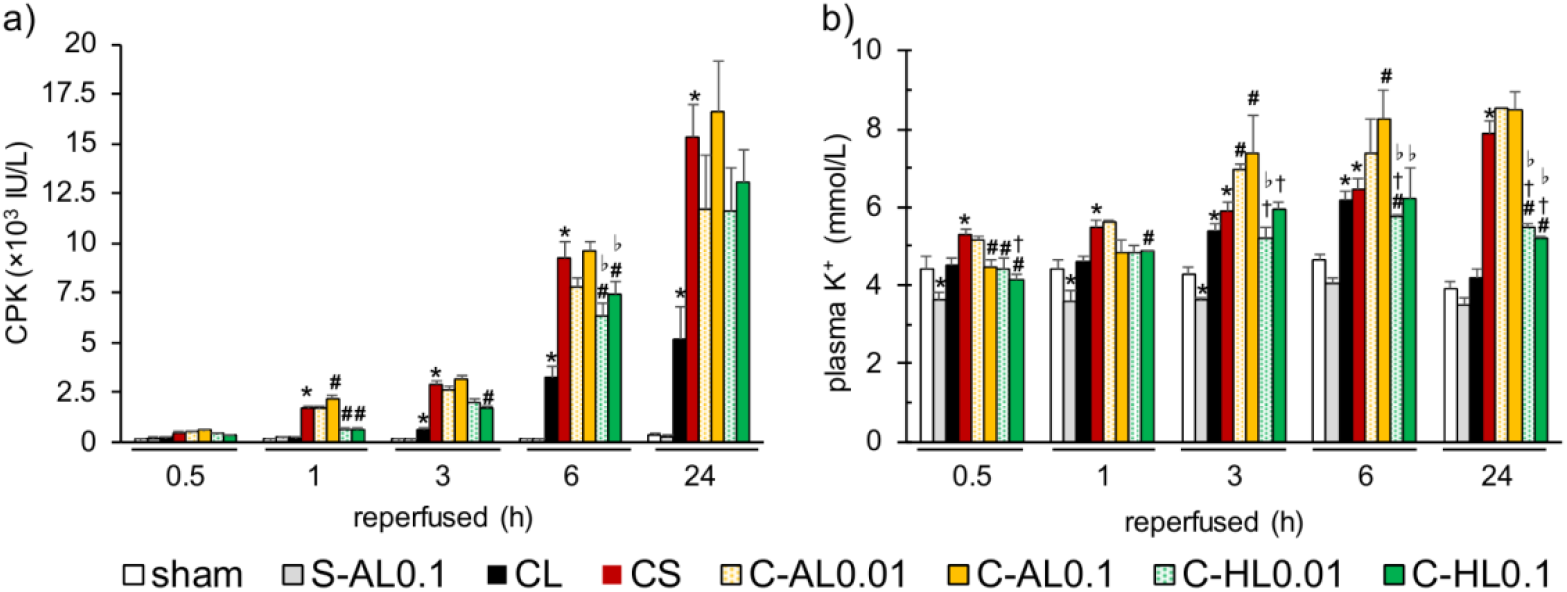
Effects of intramuscular adrenaline on cardiac failure in rats with CS. a) CPK and b) K+ levels. White bar, sham; gray bar, S-AL0.1; black bar, CL; red bar, CS; yellow dot bar, C-AL0.01; yellow bar, C-AL0.1; green dot bar, C-HL0.01; and green bar, C-HL0.1. Values represent the means±SEMs (respectively, n=6). ^*^p<0.05 vs. sham group, ^#^p<0.05 vs. CS group, ^†^p<0.05 vs. C-AL-0.01, ^♭^p<0.05 vs. C-AL-0.1, ^§^p<0.05 vs. C-HL-0.01 (Tukey–Kramer test).

Fig. 3 shows the levels of Mb, urine volume, urine pH, BUN, and creatinine, which were used to assess the effectiveness of CS against myoglobinuric acute renal failure. The Mb levels in the CS group were significantly higher than those in the sham group throughout the experimental period (513.5±57.8 vs. 5.8±1.1 mg/dL, p<0.05, reperfused at 24 h). Although the Mb levels in the C-AL0.01 and C-AL0.1 groups were similar to those in the CS group, they were significantly lower in the C-HL0.01 and C-HL0.1 groups. In the CS group, urine volume and pH significantly decreased (urine volume, 0.11±0.02 vs. 0.40±0.06 mL, p<0.05; urine pH, 4.8±0.2 vs. 6.2±0.2, p<0.05, reperfused at 24 h), whereas BUN and creatinine levels were significantly higher compared with the sham group (BUN, 125±14.6 vs. 27.3±5.49 mg/dL, p<0.05;creatinine, 1.10±0.02 vs. 0.19±0.05 mg/dL, p<0.05). In contrast, these parameters were significantly improved in the Adr-treated group. The S-AL0.1 and CL groups showed no significant differences with the sham group.

**Figure 3.**
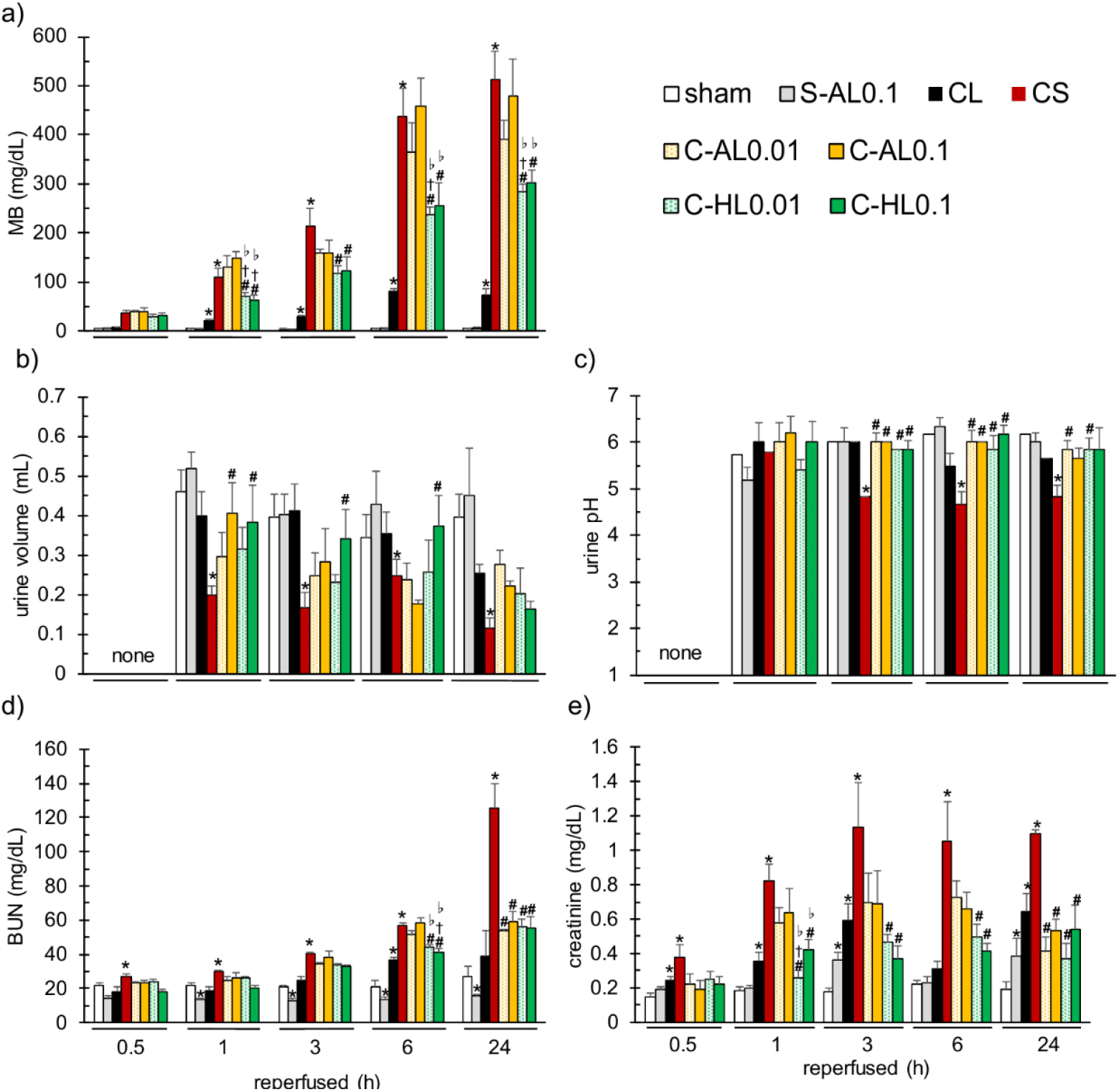
Effects of intramuscular adrenaline on AKI in rats with CS. a) MB levels, b) urine volume, c) urine pH levels, d) BUN levels, and e) creatinine levels. White bar, sham; gray bar, S-AL0.1; black bar, CL; red bar, CS; yellow dot bar, C-AL0.01; yellow bar, C-AL0.1; green dot bar, C-HL0.01; and green bar, C-HL0.1. Values represent the means±SEMs (respectively, n=6). ^*^p<0.05 vs. sham group, ^#^p<0.05 vs. CS group, ^†^p<0.05 vs. C-AL-0.01, ^♭^p<0.05 vs. C-AL-0.1 (Tukey–Kramer test).

### Effects of intramuscular adrenaline on vital signs for shock

Fig. 4 shows the SBP, HR, blood pH, BE, and BT levels. The CS group exhibited significantly lower levels of all parameters than the sham group throughout the experimental period. SBP, HR, and BT levels were significantly higher in the Adr-treated groups than in the CS group. These levels were higher in the Adr-HL group than in the Adr-AL group. In contrast, the blood pH and BE levels in the Adr-treated groups remained lower than those in the CS group throughout the experimental period. The lactate levels in the CS group were significantly higher than those in the sham group. In the C-AL0.1 and C-HL0.1 groups, lactate levels were similar to those in the CS group. The S-AL0.1 and CL groups showed no significant differences with the sham group.

**Figure 4.**
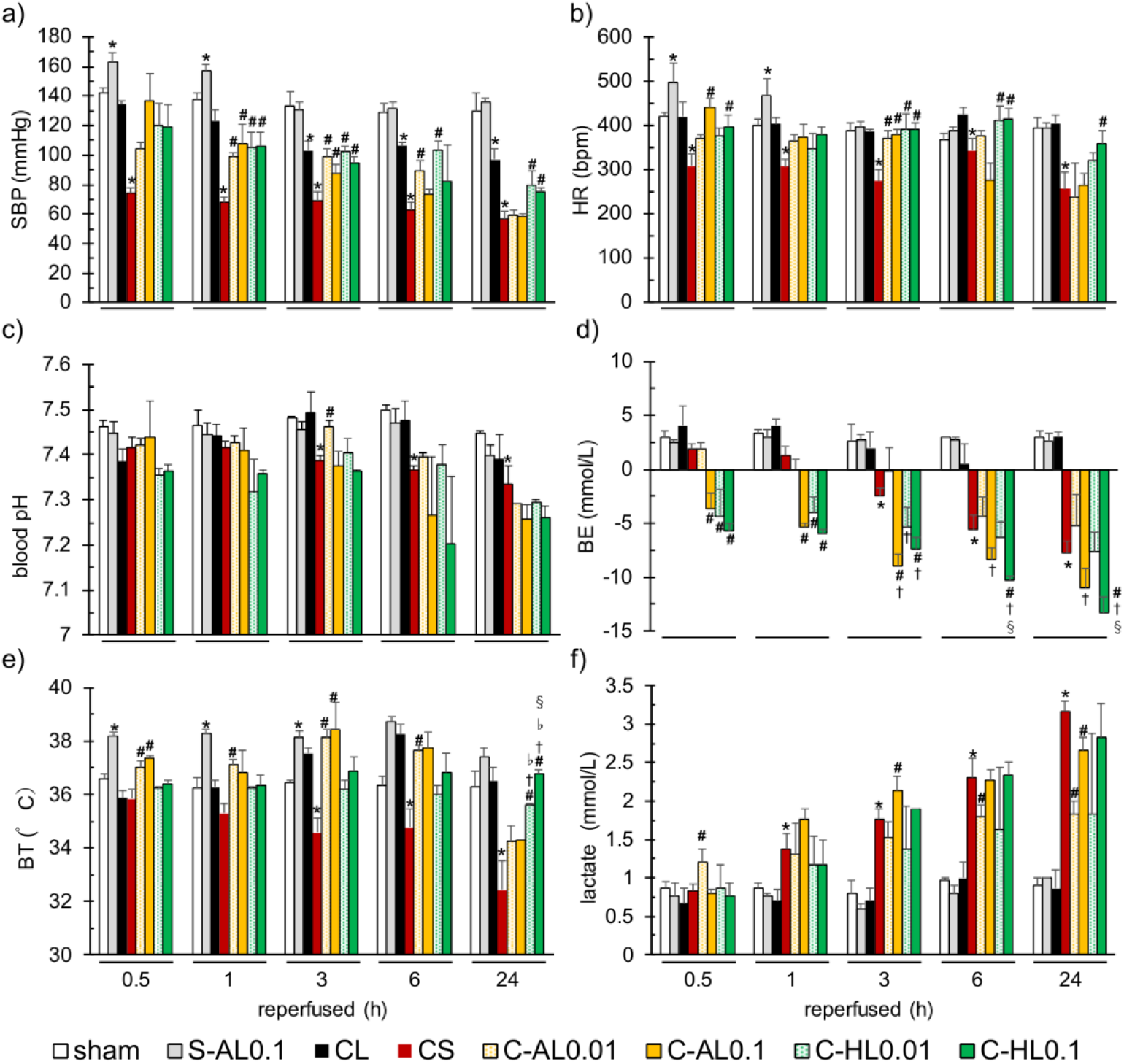
Effects of intramuscular adrenaline on shock state in rats with CS. a) SBP levels, b) HR, c) blood pH, d) BE levels, e) BT levels, and f) lactate levels. White bar, sham; gray bar, S-AL0.1; black bar, CL; red bar, CS; yellow dot bar, C-AL0.01; yellow bar, C-AL0.1; green dot bar, C-HL0.01; and green bar, C-HL0.1. Values represent the means±SEMs (respectively, n=6). ^*^p<0.05 vs. sham group, ^#^p<0.05 vs. CS group, ^†^p<0.05 vs. C-AL-0.01, ^♭^p<0.05 vs. C-AL-0.1 (Tukey–Kramer test).

### Effects of intramuscular adrenaline on injured muscle and femoral vessel blood flow

Fig. 5 shows the effects of Adr administration on blood flow in the injured muscle and femoral vessels. The blood flow levels in the S-AL0.1 group remained higher than those in the sham group throughout the experimental period. Conversely, the blood flow levels in the CL group were lower than those in the sham group during the same period. The muscle blood flow in the CS group remained significantly lower than the sham group throughout the experiment (116.6±16.0 vs. 291.8±15.0 PU, p<0.05, reperfused at 0.5 h). The Adr-AL group showed a significant increase over the CS group only at 0.5 and 1 h (174.9±6.5, 196.7±6.6 vs. 116.6±16.0 PU, p<0.05, reperfused at 0.5 h). Notably, the Adr-HL group showed lower levels than the CS group at the same time points. The femoral blood flow in the CS group exhibited a decreasing trend at 0.5 and 1 h, followed by an increasing trend at 3 and 6 h, compared with the sham group. The femoral vessel blood flow in the Adr-AL group was significantly higher compared with that in the CS group only at 0.5 and 1 h. Notably, the blood flow in the femoral vessels in the Adr-HL group was lower than that in the CS group at the same time points.

**Figure 5.**
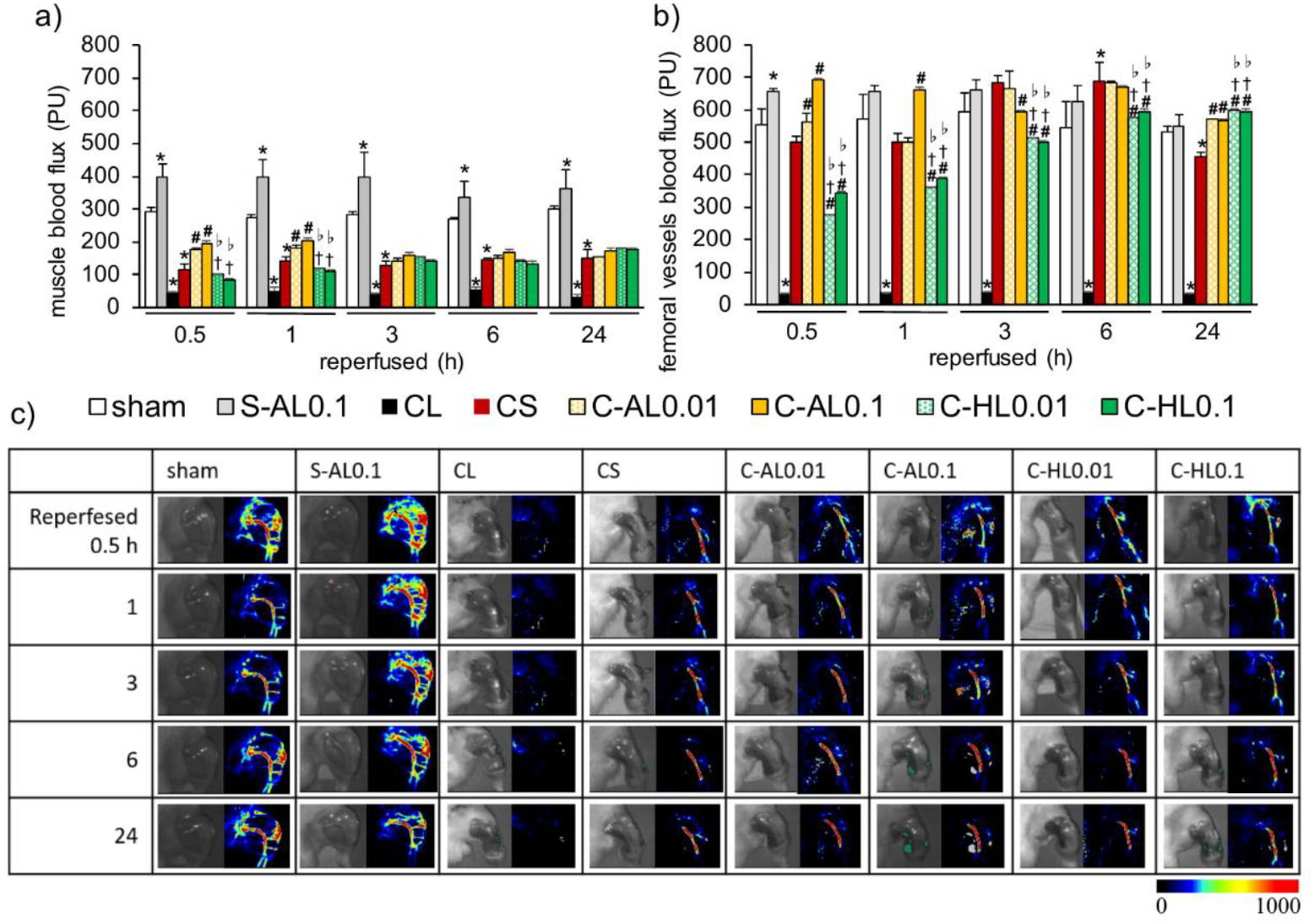
Effects of intramuscular adrenaline on blood flow in rats with CS. a) Muscle blood flow levels, b) femoral vessels blood flow levels, and c) laser Doppler image showing the changes in an affected hind limb. White bar, sham; gray bar, S-AL0.1; black bar, CL; red bar, CS; yellow dot bar, C-AL0.01; yellow bar, C-AL0.1; green dot bar, C-HL0.01; and green bar, C-HL0.1. Values represent the means±SEMs (respectively, n=6). ^*^p<0.05 vs. sham group, ^#^p<0.05 vs. CS group, ^†^p<0.05 vs. C-AL-0.01, ^♭^p<0.05 vs. C-AL-0.1, ^§^p<0.05 vs. C-HL-0.01 (Tukey–Kramer test).

### Effects of intramuscular adrenaline on inflammatory markers

Fig. 6 shows the plasma ANP and IL-6 levels. The plasma ANP and IL-6 levels were significantly higher in the CS group than in the sham group. In the C-AL0.1 group, plasma ANP levels were significantly lower than those in the CS group. In the Adr group, plasma IL-6 levels were not significantly different from those in the CS group.

**Figure 6.**
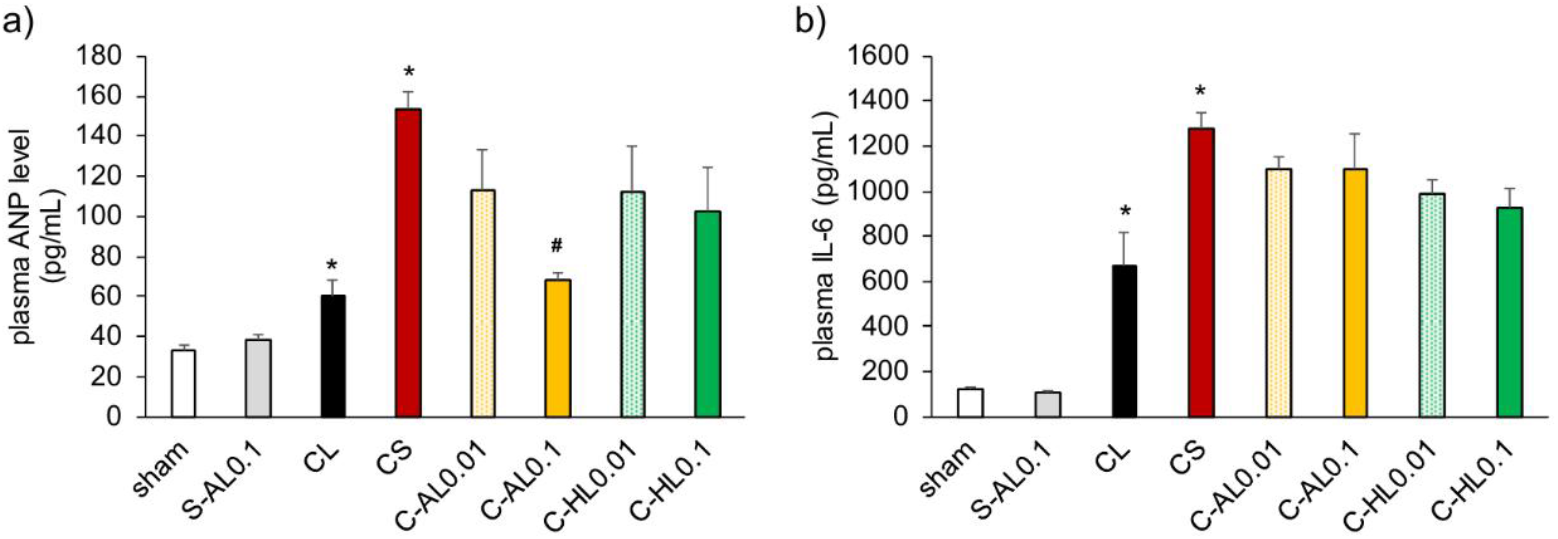
Effects of intramuscular adrenaline on cytokine in rats with CS. a) Plasma ANP and b) plasma IL-6 levels. White bar, sham; gray bar, S-AL0.1; black bar, CL; red bar, CS; yellow dot bar, C-AL0.01; yellow bar, C-AL0.1; green dot bar, C-HL0.01; and green bar, C-HL0.1. Values represent the means±SEMs (respectively, n=6). ^*^p < 0.05 vs. sham group, ^#^p < 0.05 vs. CS group (Tukey–Kramer test).

## Discussion

CS has a mortality rate of approximately 20% shortly after extrication, largely due to cardiac arrest and shock resulting from K^+^ release from injured muscles (5). In the present study, we evaluated the effectiveness of intramuscular Adr administration in a rat CS model. Our results demonstrated that although Adr significantly improved certain hemodynamic and kidney function parameters, its impact on the overall survival rate did not improve.

One of the reasons for this was hemodynamic stabilization and the K^+^ dilemma. In the untreated rats, epinephrine administration resulted in decreased K^+^ levels (Fig. 2). This effect was attributed to enhanced intracellular K^+^ uptake driven by β-adrenoceptor stimulation. Specifically, it was suggested that β1-receptor stimulation promoted renin and subsequent aldosterone secretion, whereas β2-receptor stimulation increased Na+/K+-ATPase activity, leading to a K^+^ shift from the extracellular to the intracellular space (8). However, the Adr-AL group showed a significant increase in SBP, HR, and peripheral blood flow (Fig.5, both the muscle and femur) in the early phase (0.5–1 h). This suggests that the β-adrenergic effect of Adr effectively enhances cardiac output and peripheral perfusion. Moreover, increased BT levels elevate blood K^+^ levels (9). Therefore, these changes in blood flow-promoting factors are thought to facilitate the systemic release (washout) of K^+^ from the damaged muscle tissue (Fig. 2). Thus, the high plasma K^+^ levels were attributed to AL administration, which exerted robust systemic effects at high doses. The pharmacokinetics of CS model rats show twofold higher plasma concentrations and stronger systemic effects when drugs are administered to intact muscles (i.e., the AL group) compared with injured muscles (10). In other words, unless the drug is administered directly to the injured site, the delivery volume to that area decreases by more than half, causing β-adrenergic effects to predominate over α-adrenergic effects. This “washout” of intracellular electrolytes likely counteracted the hemodynamic benefits, explaining why the AL group did not show a survival advantage. Nevertheless, Adr administration in the HL group might have shown a protective effect mediated by the suppression of the systemic circulation of nephrotoxins, such as Mb, through vasoconstriction (Fig. 3).

The Adr-HL (high-dose) group exhibited lower muscle and femoral blood flow than the CS group during the hyperacute phase. This suggests that strong β-adrenergic stimulation induces potent vasoconstriction at the site of injury (Fig. 5). Notably, this localized reduction in blood flow appeared to sequester Mb within damaged tissues, leading to significantly lower plasma Mb levels and improved renal function (Fig. 3). These findings align with those of previous studies on icing, suggesting that temporary restriction of flow from the injured limb can mitigate myoglobinuric acute renal failure (6). Despite the improvements in blood pressure and renal markers, blood pH and BE levels in all the 0.1 mg/kg Adr-treated groups remained lower than those in the CS group throughout the experimental period (Figs. 3 and 4). Although Adr stabilizes systemic circulation, this persistent metabolic acidosis may exacerbate tissue hypoxia and inhibit the clearance of lactate in the pathophysiology of CS, thereby accelerating the development of renal failure or contributing to the worsening of the condition. The fact that 48-h survival did not significantly improve in the HL group indicates that the detrimental effects of hyperkalemia and exacerbated acidemia likely surpassed the therapeutic gains from hemodynamic stabilization and renal protection. To propose the clinical significance of Adr for CS, we examined its effectiveness against inflammatory responses, a topic that remains a subject of debate. ANP suppresses the production of inflammatory cytokines (Fig. 6) (11) and provides organ-protective benefits (12,13). In contrast, Staedtke et al. reported that Adr administration promoted cytokine storms mediated by α1-receptor stimulation (7). Although inflammatory responses in CS are often masked by acute-phase symptoms, patients frequently progress to inflammatory conditions leading to death after heart or acute kidney failure (14). Therefore, managing inflammation is a critical strategy in the treatment of CS. However, Adr administration did not induce inflammation in the present study. The significant reduction in plasma ANP levels in the Adr-treated groups suggests a reduction in cardiac wall stress, possibly owing to stabilized hemodynamics.

Although a rat model of CS provides valuable physiological insights, it may not perfectly replicate the complex clinical course of CS in humans. Specifically, the “washout” effect of K^+^ from injured muscle observed in our model might occur differently in human patients depending on the scale of injury and the timing of fluid resuscitation. We concluded that Adr did not induce a cytokine storm. However, our observation period was limited to the acute phase (up to 48 h). As patients with CS often progress to fatal inflammatory conditions after surviving the initial heart or kidney failure, a longer observation period might be necessary to fully evaluate the long-term immunomodulatory effects of Adr.

In conclusion, intramuscular Adr provides immediate hemodynamic stabilization and renal protection during the early stages of CS. However, its use requires careful dose titration; low doses may promote the systemic release of lethal toxins, whereas high doses may worsen metabolic acidosis. These findings highlight the importance of combining Adr with other therapies, such as fluid resuscitation, to manage systemic toxemia inherent in CS.

## Abbreviations

CS: crush syndrome
K+: potassium
MB: myoglobin
Adr: adrenaline
AL: anterior limb
HL: hind limb
CL: clamping
SBP: systolic blood pressure
MAP: mean arterial pressure
DBP: diastolic blood pressure
HR: heart rate
BT: body temperature
E: base excess
BUN: blood urea nitrogen
CPK: creatine phosphokinase
IL: interleukin
ANP: atrial natriuretic peptide
BPF: blood perfusion
SEM: standard error of the mean.

## Acknowledgments

The authors thank Ms. Marise Komiya, Ms. Shion Terada, Ms. Yumiko Murakawa, and Ms. Chikako Murata for their technical assistance.

## Availability of data and materials

Not applicable

## Authors’ contributions

IM led the project and designed and performed most of the experiments. IM and YM assisted with the survival and biochemical marker analyses. JK and YM conceived the study, participated in its design and coordination, and drafted the manuscript. All the authors have read and approved the final manuscript.

## Ethics approval

All animal experiments were approved by the Institutional Animal Care and Use Committee of Josai University (approval nos.: H29030 and JU18031).

## Patient consent for publication

Not applicable

## Authors’ information

## References

1. Demirkiran O, Dikmen Y, Utku T and Urkmez S: Crush syndrome patients after the Marmara earthquake. Emerg Med J ITR, Kennedy 20: 247–250, 2003, Kennedy ITR, Petley DN, Williams R and Murray V: A Systematic review of the health impacts of mass earth movements (landslides). PLOS Curr 7: ecurrents.dis.1d49e84c8bbe678b0e70cf7fc35d0b77, 2015.

2. Usuda D, Shimozawa S, Takami H, Kako Y, Sakamoto T, Shimazaki J, Inoue J, Nakayama S, Koido Y and Oba J: Crush syndrome: a review for prehospital providers and emergency clinicians. J Transl Med 21: 584, 2023.

3. Gonzalez D: Crush syndrome. Crit Care Med 33 Suppl: S34–S41, 2005.

4. Murata I, Ooi K, Sasaki H, Kimura S, Ohtake K, Ueda H, Uchida H, Yasui N, Tsutsui Y, Yoshizawa N, et al.: Characterization of systemic and histologic injury after crush syndrome and intervals of reperfusion in a small animal model. J Trauma 70: 1453–1463, 2011.

5. Ashkenazi I, Isakovich B, Kluger Y, Alfici R, Kessel B and Better OS: Prehos. Disast.Med. 20: 122–133, 2005.

6. Murata I, Imanari M, Komiya M, Kobayashi J, Inoue Y and Kanamoto I: Icing treatment in rats with crush syndrome can improve survival through reduction of potassium concentration and mitochondrial function disorder effect. Exp Ther Med 19: 777–785. doi: 10.3892/etm.2019.8230[ePub]2019 Nov 22, 2020.

7. Staedtke V, Bai RY, Kim K, Darvas M, Davila ML, Riggins GJ, Rothman PB, Papadopoulos N, Kinzler KW, Vogelstein B, et al.: Disruption of a self-amplifying catecholamine loop reduces cytokine release syndrome. Nature volume 564, pp 273–277, 2018.

8. Haalboom JRE, Deenstra M and Struyvenberg A: Hypokalaemia induced by inhalation of fenoterol. Lancet 1: 1125–1127, 1985, Wang P and Clausen T: Treatment of attacks in hyperkalaemic familial periodic paralysis by inhalation of salbutamol. Lancet 1: 221–223, 1976.

9. Nakayama T and JSR, 188: 250–259, 2014.

10. Murata I, Goto M, Komiya M, Motohashi R, Hirata M, Inoue Y and Kanamoto I: Early therapeutic intervention for crush syndrome: characterization of intramuscular administration of dexamethasone by pharmacokinetic and biochemical parameters in rats. Biol Pharm Bull 39: 1424–1431, 2016.

11. Morita R, Ukyo N, Furuya M, Uchiyama T and Hori T: Atrial natriuretic peptide polarizes human dendritic cells toward a Th2-promoting phenotype through its receptor guanylyl cyclase-coupled receptor A. J Immunol 170: 5869–5875, 2003.

12. Kitakaze M, Asakura M, Kim J, Shintani Y, Asanuma H, Hamasaki T, Seguchi O, Myoishi M, Minamino T, Ohara T, et al.: Human atrial natriuretic peptide and nicorandil as adjuncts to reperfusion treatment for acute myocardial infarction (J-WIND): two randomised trials. Lancet 370: 1483–1493, 2007.

13. Kasama S, Furuya M, Toyama T, Ichikawa S and Kurabayashi M: Effect of atrial natriuretic peptide on left ventricular remodelling in patients with acute myocardial infarction. Eur Heart J 29: 1485–1494, 2008.

14. Zhang H, Zeng JW, Wang GL, Tu CQ, Huang FG and Pei FX: Infectious complications in patients with crush syndrome following the Wenchuan earthquake. Chin J Traumatol 16: 10–15, 2013.

